# *In situ* humoral selection in human lupus tubulointerstitial inflammation

**DOI:** 10.1101/2020.06.29.178145

**Authors:** Andrew J Kinloch, Yuta Asano, Azam Mohsin, Carole J Henry, Rebecca Abraham, Anthony Chang, Christine Labno, Patrick C Wilson, Marcus R Clark

## Abstract

In human lupus nephritis, tubulointerstitial inflammation (TII) is associated with *in situ* expansion of B cells expressing anti-vimentin antibodies (AVAs). The mechanism by which AVAs are selected is unclear. Herein, we demonstrate that AVA somatic hypermutation and selection increase affinity for vimentin. However, enzyme-linked immunosorbent assays (ELISAs) suggested that affinity maturation might be a non-specific consequence of increasing polyreactivity. Subsequent multi-color confocal microscopy studies indicated that while TII AVAs often appeared polyreactive by ELISA, they bound selectively to vimentin fibrils in whole cells or inflamed renal tissue. Using a novel machine learning pipeline (CytoSkaler) to quantify the cellular distribution of antibody staining, we demonstrated that TII AVAs were selected for both enhanced binding and specificity *in situ*. These data suggest a new approach to assess and define antibody polyreactivity based on quantifying the distribution of binding to native and contextually relevant antigens.

## INTRODUCTION

Nephritis is the most common severe manifestation of Systemic Lupus Erythematosus (SLE). Lupus nephritis canonically is thought of as a glomerulonephritis (GN) with extensive evidence in both humans and mice indicating that lupus GN is a manifestation of systemic autoimmunity [1]. However, inflammation and scarring in the tubulointerstitium, and not in glomeruli, predict progression to renal failure [2, 3]. Furthermore, tubulointerstitial inflammation (TII) is associated with complex local adaptive immunity including *in situ* B cell selection [4, 5] and help provided by resident T follicular helper-like cells [6]. These findings suggest that the immunological mechanisms that drive GN and TII in lupus are very different.

Previously, we have isolated antibodies expressed by clonally expanded B cells in human lupus TII [7]. Remarkably, of 25 monoclonal antibodies (mAbs) cloned from eight patients, ten from six of these patients directly bound vimentin. In a cross-sectional cohort, high serum anti-vimentin antibodies (AVAs) correlated with severe TII on renal biopsy [7]. Finally, in the LUNAR trial of Rituximab in lupus nephritis, high serum AVAs at study entry predicted a poor outcome regardless of treatment arm [8]. These data suggest that AVAs are a feature of severe TII that predicts progressive lupus renal disease.

Vimentin is strongly upregulated in renal inflammation being expressed both by infiltrating T cells and macrophages as well as stressed tubulo-epithelium [7, 9, 10]. Therefore, our studies suggest that in lupus nephritis, immune tolerance can be broken *in situ* to molecular patterns of inflammation and damage. This is in contrast to typical lupus peripheral blood antibody specificities that target nucleotide-protein complexes [11]. Elegant studies have demonstrated that these latter peripheral specificities are associated with somatic hypermutation and selection for high affinity [12-14]. However, in many cases, somatically hypermutated and selected autoantibodies, such as those that target dsDNA, are reactive to a broad range of antigens *in vitro* [15, 16]. These studies suggest that in lupus, autoantibodies are selected for affinity but not necessarily for specificity.

Herein, we demonstrate that AVA somatic hypermutation confers both increased vimentin affinity and broad polyreactivity in *in vitro* assays. However, when we used confocal microscopy and machine learning to quantify *in situ* binding, it was apparent that polyreactivity as measured by ELISA, correlated with both increased binding and specificity for vimentin fibrils. These results suggest that ELISA antibody reactivities do not predict *in vivo* binding specificity.

## MATERIALS AND METHODS

### Study Subjects

Patients from which TII antibodies were cloned have been described previously [7]. For staining lupus nephritis biopsies, paraffin embedded kidney diagnostic biopsies from patients meeting ACR criteria for lupus and having severe TII (as confirmed by renal pathologist AC) were used. Samples were de-identified and used under IRB 15065B.

### Reagentss

Commercial antigens were double stranded DNA (cat. D3664, SIGMA), insulin (I9278, SIGMA) and LPS (L6143, SIGMA). Confocal related reagents included antigen retrieval buffer low pH (cat. 00-4955-58, Invitrogen), donkey serum (cat. 017-000-121, Jackson ImmunoResearch), human Fc block (cat.584220, BD Phramingen) and Hoechst 33342 (cat.H3570, ThermoFisher), ProLong Gold anti-fade mounting solution (P36930, ThermoFisher). Commercial primary antibodies included, murine IgG anti-vimentin (clone V9, DAKO), rabbit IgG anti-enolase 1 (clone EPR 10863, Abcam) and rat IgG anti-DYKDDDDK (anti-FLAG, clone L5, Biolegend). Secondary antibodies (ThermoFisher) were Alexa Fluor 488 goat anti-rat IgG, Alexa Fluor 647 donkey anti-rabbit IgG, Alexa Fluor 568 anti-mouse IgG 568, Alexa Fluor 488 goat anti-human IgG and anti-human IgG HRP (Jackson ImmunoResearch).

### Cloning and antibody purification

Human IgG1 mAbs were cloned and expressed as reported previously [7, 17]. Germline reverted antibody clones were generated by expressing pairs of corresponding heavy and light chain expression constructs encoding somatic hypermutations altered to their predicted germline sequences using IMGT [18]. For the TII mAb PB5, cDNAs encoding each individual immunoglobulin heavy chain somatic hypermutation reversion were likewise generated and respectively paired with fully somatic hypermutated immunoglobulin light chains. All reverted variable region nucleotide sequences were purchased as synthetic sequences (Genscript) and subcloned into the respective heavy or light chain expression vectors. Full-length vimentin in vector pET-24b [10] was used for *in vitro* expression (9BL21 Rosetta DE3 pLysS, EMD) and purification [19]. For performing tissue staining, a novel FLAG tagged IgG1 fusion protein was engineered. The Igg chain expression vector FJ475055, was modified by inserting the nucleotide sequence (GACTACAAAGACGATGACGACAAG) immediately 5’ to the stop codon.

Following co-expression with the light chain plasmid, the recombinant human IgG1 harbored C-terminal FLAG tags on each heavy chain, enabling purification with protein A as performed previously [7, 17].

### ELISA

Anti-vimentin antibody ELISAs were performed by diluting antigen to 10 μg/ml in PBS before coating wells of ELISA plates (cat.no 3690, Corning Costar). Control wells were coated with blocking buffer (PBS/3% BSA). Plates were rinsed with ddH20 and blocked (PBS/3% BSA) for two hours. Primary antibodies diluted (30 μg/ml) in PBS were incubated for two hours and washed three times with PBS/Tween 0.01%. Bound IgG was detected using HRP-conjugated secondary antibodies diluted 1:1000 in blocking buffer. Following washing (as for the primary incubation), Super AquaBlue (eBioscience) was added and optical density measured at 405 nm. Calculation of *Kds* was performed using Graphpad Prism following a succession of three-fold dilutions as reported previously [20]. Commercial assays were used for assaying anti-histone (QUANTA Lite 708520, Inova Diagnostics) and ant-dsDNA (QUANTA Lite 708510, Inova Diagnostics) antibody reactivity in Supplemental Figure 2 according to manufacturer’s protocols.

Polyreactivity ELISAs were performed as reported previously [15, 16]. Plates were incubated at 4 °C overnight with carbonate buffer (0.15 M Na2CO3, 0.35 M NaHCO3) alone or carbonate buffer containing either 10 μg/mL dsDNA, 10 μg/mL LPS, or 5 μg/mL insulin. Plates were washed 3 times with ddH2O and treated with 1 mM EDTA/0.05% Tween-PBS at 37 °C for 1 hour. Plates were washed 3 times with ddH2O and incubated with 1 μg/mL antibodies at 37 °C for 1 hour. Plates were washed with ddH2O and incubated with HRP-conjugated 2^nd^ antibody diluted 1:2000 in 1 mM EDTA/0.05% Tween-PBS at 37 °C for 1 hour. After 3 wash with ddH2O, plates were incubated with 1mM EDTA/0.05% Tween-PBS at room temperature for 5 minutes, and washed 3 times with ddH2O. Signal was developed by Super AquaBlue and measured at 405 nm.

### Cell Staining

For multi-color HEp-2 cell imaging, cells were cultured (10,000 cells per well) on poly-L-lysine five well Teflon-coated slides (cat.185-051-120, TEKDON) for 20 hours, fixed in methanol, and stained as previously reported [7]. Primary antibodies were human IgG mAbs (50 μg/ml), V9 and anti-Eno-1 (1μg/ml). Fluorophore conjugated secondary antibodies (listed above,1:1000) and Hoechst (1:500) were added after washing. After a second round of washing, pro-long gold mounting medium was added to each well, cover slips were attached and slides were kept in the dark until imaging.

### Tissue staining

Slides from paraffin embedded lupus kidney biopsies were prepared and stained as previously described [19]. Primary antibodies were added at the concentrations indicated for HEp-2 cell analyses. Rat IgG anti-DYKDDDDK (1:200) was used as a secondary antibody to detect tagged human IgG mAbs. Tertiary antibodies, Alexa Fluor 488 anti-rat IgG and Alexa Fluor 568 anti-mouse IgG, were added concomitantly with Hoechst (1:500). After a final round of washing pro-long gold mounting medium was added and cover slips attached.

### Image Acquisition

Slides were analyzed on a Leica SP8 Leica STED-CW laser scanning confocal microscope using the oil immersion objective (63x). Fields of view were 1024 x 1024 pixels at 16-bit depth using a white light laser. Respective stains were acquired in separate channels. For any given session in which human mAbs were being compared quantitatively, photomultiplier tube (PMT) and objective magnifications were constant. Channels were saved as separate raw TIF files. To process images, TIF files for respective channels were assigned in Image J freeware and merged.

### Manual image analyses

The boundaries of HEp-2 cells were defined with the Image J polygon free hand drawing tool. Nuclear (“nuc”) regions and areas high in vimentin (Vim^hi^) were defined using Moments and Otsu autothresholding on Hoechst and V9 channel derived raw TIFs respectively, in Image J. The mean pixel intensity (MPI) and pixel area data were acquired for each whole cell and respective Vim^hi^ and nuc regions. Manually segmented Vimentin low “Vim^lo^” area was defined as that cytoplasmic area not attributable to the Vim^hi^ zone. Cytoplasm was defined as that area in manually segment cells not attributable to the nucleus. Prior to CytoSkaler analyses (**Fig.1-2**), the JACoP plugin was used to calculate the Pearson correlation coefficients between intensities of matched pixels for whole fields of view (FOVs) in indicated channels [21].

**Figure 1.**
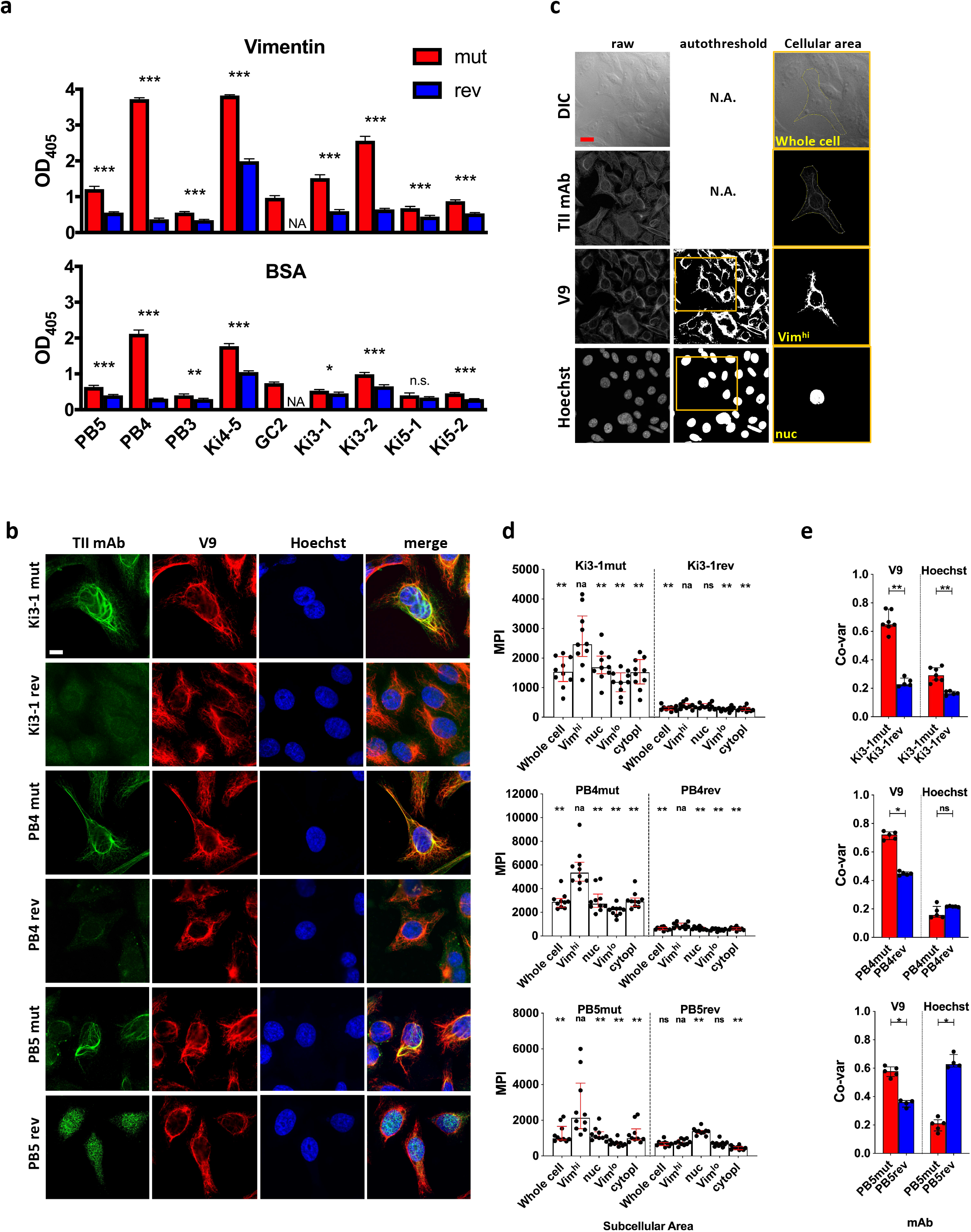
Antigen binding of germline reverted TII AVAs. Nine somatically hypermuted (mut) AVAs and their corresponding predicted germline reversions (rev) were assayed for autoreactivity. (**a**) Reactivity with recombinant vimentin or BSA were measured by ELISA (Raw OD405 values are given and t-tests were performed between mut and rev variants of respective AVAs). (**b**) Representative HEp-2 multi-color immunofluorescence microscopy images stained with indicated mutated and reverted TII antibodies with vimentin (V9) and nucleus (Hoechst). Representative of 3 independent experiments. (**c**) HEp-2 multi-color immunofluorescence microscopy method for quantification of relative AVA reactivity with different subcellular areas. Individual raw channels including DIC were used for segmenting whole cells and subcellular areas. Areas high in reactivity with Hoechst and V9 were autothresholded and designated nuclear (“nuc”) and vimentin^high^ (Vim^hi^) respectively. Cell perimeters were manually segmented. Representative example provided. (**d**) Relative reactivities (measured as mean pixel intensities, MPI) of AVAs with different subcellular areas. Each dot represents the MPI in the AVA channel for the specified subcellular zone of an individual cell. Pairwise comparisons (Wilcoxon matched-pairs tests) were performed between MPIs of Vim^hi^ and indicated subcellular areas. (**e**) Co-variances of pixel intensities were quantified between the AVA channel and either V9 or Hoechst channels. Dots represent correlation coefficients between pixel intensities for multicellular fields of view (FOV). Mann-Whitney tests were performed between somatically hypermutated (mut) and germline reversion values. For 1d, binding in each group was compared to vimentin binding. na = not applicable; ns = not statistically significant. *p < 0.05, **p < 0.01, ***p < 0.001, ****p < 0.0001. Red scale bar = 10 microns.

**Figure 2.**
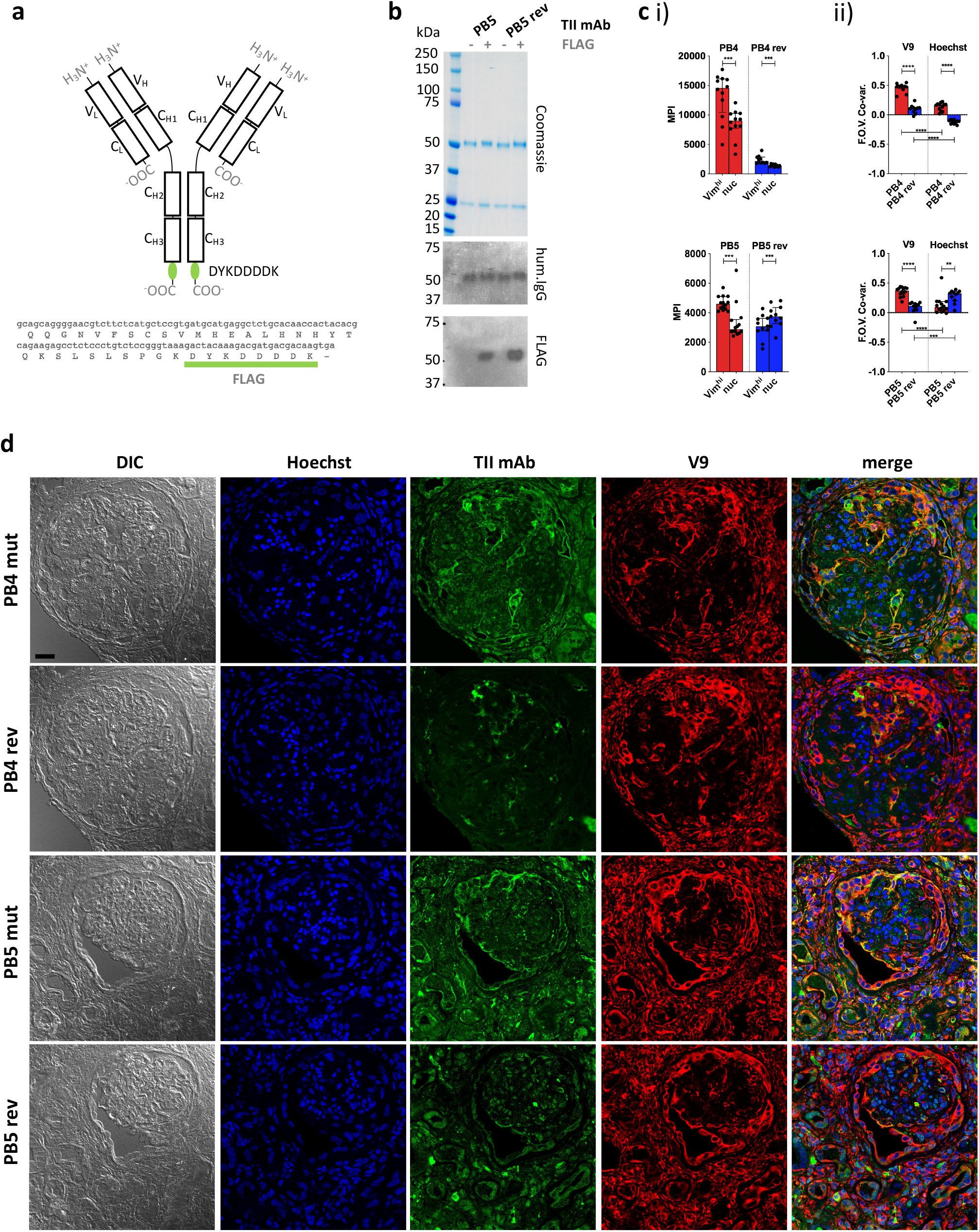
Quantitative staining of IgG rich lupus kidney. (**a**) The human IgG1 mAb heavy chain vector was genetically modified to express the FLAG sequence (DYKDDDDK) at the C-terminus of each heavy chain, enabling detection with an anti-FLAG secondary antibody. Top = Protein schematic of respective domains of IgG1 with inserted tag in green. Bottom = C-terminal amino acid and 3’ nucleotide sequences of heavy chain construct. (**b**) SDS PAGE and subsequent Coomassie staining or western blotting with anti-human IgG (hum.IgG) or anti-FLAG IgG on the selected and reverted PB5 TII mAb in the absence or incorporation of the FLAG tag. (**c**) Reactivity of FLAG-tagged PB4 (top) and PB5 (bottom) variants with Vim^hi^ and nuclear zones of inflamed lupus kidney as determined by multi-color confocal microscopy. (**i**) Each dot represents the MPI of the TII mAb channel for the indicated subcellular region for a respective FOV (**ii**) Co-variances of MPIs between the AVA and V9 or Hoechst channel. Dots represent correlation coefficients between the TII mAb channel and V9 or Hoechst channel for respective FOVs. Wilcoxon matched-pairs tests were performed between MPIs of Vim^hi^ and nuc subcellular areas. Mann-Whitney tests were performed between somatically hypermutated (mut) and germline reversion values. na = not applicable; ns = not statistically significant. *p < 0.05, **p < 0.001, ***p < 0.001, ****p < 0.0001. (**d**) Representative examples of co-staining of lupus renal tissue with FLAG-tagged PB4 and PB5 AVA variants. Black scale bar = 25 microns.

Kidney tissue was analysed as for cultured HEp-2 cells, with the exception that Vim^hi^ and nuclear regions were applied such that each was an entire FOV. Therefore, for each FOV there was one Vim^hi^ and one nuclear region. The AVA MPIs were plotted for the indicated subcellular regions for each FOV.

CytoSkaler was developed to enable automated cellular and subcellular segmentation, and subsequent quantification of TII mAb binding within each respective cell. All image analysis was conducted using MATLAB R2019a (Natick, USA) for CytoSkaler development. TIFF files from 63x magnification multicellular FOVs included raw Hoechst, V9, and anti-ENO1 stains. Images were thresholded using Otsu, Moments, and Percentile thresholding functions respectively (functions were imported from Image-J as a Java Class Object). Binary anti-ENO1 images also underwent one round of despeckling and space filling to incorporate weakly stained cell areas. Binary objects from the Hoechst stain were overlaid onto the V9 and ENO1 stains resulting in composite RGB images. Each composite RGB image contained a single Hoechst binary object overlaid in a green color (RGB binary value: 0 1 0) while the other Hoechst binary objects were overlaid in a red color (RGB: 1 0 0).

Two DeepLab v3+ convolutional neural networks (CNN’s) were trained to recognize subcellular pixels around green Hoechst binary objects for both the V9 and anti-ENO1 RGB images, using weights initialized from a pre-trained Resnet-18. Two separate networks were developed because pixel patterns were different in the V9 and anti-ENO1 channels. Each network was trained to classify each pixel into 1 of 5 classes (1. subcellular pixels of interest to the local green Hoechst boundary, 2. green Hoechst boundary, 3. other subcellular pixels near red Hoechst boundaries, 4. red Hoechst boundaries, 5. black background). Class weights for the Pixel Classification Layer were balanced using inverse pixel frequency.

The ground truth data included 1024×1024 RGB images and corresponding categorical images with class labels for each pixel (manually assigned with MATLAB Image Labeler). For each network (Vim^hi^ and Cytoplasmic), 189 images were used for training, 24 images were used for validation, and 23 images were used for testing. An image data augmenter was created to configure a set of pre-processing options that expanded the size of the training set by creating new images with variations in image rotation, isotropic scale, x-axis reflection, and y-axis reflection, during training. Ground truth data was created by manually selecting subcellular boundaries of each cell using Image-J.

Stochastic gradient descent was used for optimization, with a momentum value of 0.9, a learning rate of 1e-3, a learning rate drop period of 690 iterations, a learning rate drop factor of 0.3, and a piecewise learning rate schedule. Both networks were trained for 2300 iterations. The performance of the model was measured by cross-entropy loss. Mean IOU’s were calculated after a post-processing algorithm.

Built-in Matlab functions (including “mean” and “corrcoef”) were used to yield outputs of mean pixel intensity, subcellular area distribution, and correlation coefficients when quantifying AVA binding within each respective cell (and/or subcellular zone) after two CNN’s segmented whole cells from multicellular images. Ground truth datasets can be found at: https://github.com/awezmm/CytoSkaler.data/tree/master/data. The DAGNetwork objects can be found at: https://github.com/awezmm/CytoSkaler.data/tree/master/networks.

The graphical user interface was developed using MATLAB App Designer. The interface allows for the analysis of any custom non-overlapping zone within a single cell, in addition to segmentation of multicellular images. While machine learning models were only trained to segment Cytoskeleton-like and Cytoplasmic-like zones, the interface also allows for integration of other neural networks and thresholding algorithms for various subcellular regions.

### Statistical analysis

GraphPad Prism v.7 software (GraphPad Software, Inc., La Jolla, CA, USA) was used for graph production and calculations of statistical differences using both manual and CytoSkaler derived datasets.

### Comparison against CellProfiler

A set of 23 images, that were not part of the CytoSkaler training set, were used to generate a comparison of IOU scores in the segmentation of cytoskeleton (Vim) and cytoplasm (ENO-1) subcellular areas. Multiple CellProfiler pipelines were tested and the one with the highest IOU scores was chosen for the final comparison. The chosen pipeline involved the *IdentifyPrimaryObjects* module with min and max object diameter as 90 and 150, and the *IdentifySecondaryObject* module with ‘propagation’ as the method to identify secondary objects, ‘global’ as the threshold strategy, ‘minimum cross entropy’ as the thresholding method, 0.0 for the thresholding smoothing scale, 1.0 for the thresholding correction factor, 0.0 and 1.0 for the lower and upper bounds on threshold, and 0.05 as the regularization factor. A binary Hoechst channel image was used as the input for the *IdentifyPrimaryObjects* module and grayscale V9 and anti-ENO-1 were used as the inputs to the *IdentifySecondaryObject* module. IOU scores for CellProfiler and CytoSkaler were determined by comparing each tool’s output to manually segmented subcellular regions.

## RESULTS and DISCUSSION

We sought to understand the mechanisms governing selection of TII-associated AVAs in lupus nephritis. As reported previously [7], these antibodies are highly somatically mutated with an average of 14 mutations in the immunoglobulin heavy chain variable region (range: 8-23) and 11 mutations in the immunoglobulin light chain (range: 0-23). For nine TII AVAs [7], we reverted DNA mutations for both the heavy and light immunoglobulin chains to encode predicted amino acid germline sequences (**Supplemental Table 1**). We then expressed the reverted (rev) antibodies, and their respective somatically hypermutated (mut) versions, and studied their *in vitro* immunoreactivities by ELISA.

As reported [7], all nine mut AVAs had demonstrable binding to vimentin (**Figure 1a**). Some antibodies, such as PB4 and Ki4-5, bound vimentin strongly while some, such as PB5 and Ki5-2, bound more weakly. Reversion of the AVAs significantly diminished binding of all eight antibodies that could be expressed.

Estimation of vimentin binding affinities for several antibodies (PB4, PB5, Ki1-1, GC2, Ki3-1, Ki5-2) revealed moderate to high affinities ranging from a Kd of 7.2×10^−9^ to 7.9×10^−7^ (**Supplemental Table 2**). It was not possible to calculate affinities for most of the reversions (data not shown). These data suggest that TII AVAs undergo canonical affinity maturation.

Interestingly, the mut AVAs also displayed some immunoreactivity with the blocking reagent used in the above assay, bovine serum albumin (BSA) (**Figure 1a**). Furthermore, binding was generally proportional to that binding ascribed to vimentin immunoreactivity. These results suggest that high vimentin binding might be associated with broad immunoreactivity. Therefore, we next assayed binding to a panel of antigens commonly used to assess polyreactivity including double-stranded (ds) DNA, insulin and lipopolysaccharide (LPS) (**Supplemental Figure 1**) [15, 16]. Indeed, most of the antibodies, except for PB5 and PB3, appeared to bind strongly to all three substrates in the polyreactive antigen panel suggesting broad reactivity. As was observed for vimentin binding, reversion to germline greatly diminished apparent polyreactivity for all antibodies except Ki4-5. The antibody Ki4-5 had strong broad binding that was only moderately diminished by reversion to germline.

Usually, the above assays of polyreactivity are done without a blocking reagent [15, 16, 22]. Therefore, as a control, we assayed AVA binding to ELISA plate wells without antigen. Surprisingly, a similar pattern of binding was observed with some mut antibodies demonstrating high binding to the uncoated plate. From these ELISA results, it is unclear if there is *in vivo* selection for vimentin binding affinity, selection for broad immunoreactivity or just acquisition of non-specific “stickiness”.

These ELISA binding characteristics were in marked contrast to what was observed when HEp-2 cells were stained with a sample of three TII AVAs (**Figure 1b**) [7]. As demonstrated, staining with Ki3-1mut, PB4mut and PB5mut provided a fibrillar pattern of cytoplasmic staining that extensively overlapped with the distribution of vimentin fibrils [23]. In general, the AVAs appeared to bind to a subset of vimentin fibrils suggesting specificity for particular isoforms or conformations. With all three antibodies there was comparatively little immunoreactivity with either the nucleus or cytoplasm not containing vimentin.

Reversion of both Ki3-1mut and PB4mut to germline resulted in a general loss of HEp-2 immunoreactivity. In contrast, PB5rev bound strongly to the nucleus. This was not associated with significant binding to dsDNA or any other polyreactive substrates by ELISA.

Both ELISA binding and staining of HEp-2 cells demonstrated that somatic hypermutation was associated with increased binding strength to vimentin. Most reverted antibodies had both poor binding to substrates by ELISA and by HEp-2 cell confocal analyses suggesting selection from low affinity clones. In contrast, PB5rev had readily apparent nuclear binding indicating selection from a previously autoreactive clone with subsequent loss of that autoreactivity and acquisition of vimentin binding. These data suggest that selection for vimentin reactivity can arise from both autoreactive and non-autoreactive naïve precursor B cells.

ELISA and confocal studies revealed remarkably different results in regard to specificity. By ELISA, increased affinity for vimentin was usually associated with increased broad, non-specific binding. In contrast, by confocal microscopy, increased somatic hypermutation and vimentin binding appeared associated with increased specificity and, in the case of PB5mut, loss of other binding specificities. Assaying *in vitro* binding to a purified antigen is artificial in that *in vivo* binding is the end-product of competition between thousands of potential antigenic epitopes. Furthermore, with purified antigen-based ELISAs, biological context, and possibly critical conformational features, are lost. As demonstrated, for some antibodies assessing binding by ELISA can be very misleading. Therefore, we developed a quantitative approach to assess relative binding strength and specificity in a cellular context.

For these initial studies, we determined the relative binding intensities of TII AVAs to different structures in the cell. To define vimentin rich zones, V9 staining (**Figure 1c**) was used. As a nuclear marker we used Hoechst while the extent of the cytoplasm was defined by manual segmentation of differential interference contrast (DIC) images. Moments and Otsu autothresholding were used on the V9 and Hoechst channels respectively to define the pixel locations of vimentin high (Vim^hi^) and nuclear (nuc) zones. Preferential binding of AVAs to respective compartments could therefore be measured by comparing antibody binding, as measured by mean pixel intensity (MPI), across the different subcellular zones. As a second metric for antibody binding, pixel intensity correlations were calculated between the test antibody channel and either the V9 or Hoechst channels.

We first examined the MPI of binding across the cellular compartments for mutated and reverted forms of Ki3-1, PB4 and PB5. As demonstrated (**Figure 1d**), all three AVAs preferentially bound the Vim^hi^ cytoplasmic compartment. Conversely, all three antibody reversions had greatly diminished binding across all three subcellular compartments, except for PB5rev, which had relatively increased binding to the nucleus. The increased magnitude of Vim^hi^ binding with somatic hypermutation was predicted by ELISA. However, only quantitative confocal microscopy revealed that increasing binding was associated with increased specificity.

By applying co-variance analyses to MPIs between AVA and either V9 or Hoechst channels, an even clearer picture became apparent (**Figure 1e**). Reversion of Ki3-1mut resulted in a loss of colocalization with V9. A similar decrease was observed for PB4mut. Interestingly, there was also a decrease in colocalization with Hoechst for Ki3-1rev and slight increase for PB4rev. However, the magnitude of these differences was much less than those observed for vimentin colocalization. In contrast, reversion of PB5mut was associated with a strong increase in co-variance with Hoechst and decreased co-variance with V9. From these experiments, we conclude that quantitative imaging captures the binding specificities of TII-associated AVAs.

We next sought to further examine *in situ* specificity by directly assessing TII AVA binding to lupus nephritis biopsies. In lupus nephritis, IgG expressing B-cells and immune complexes are wide-spread. Therefore, to avoid detection of endogenous IgG with anti-human IgG secondary antibodies, we epitope-tagged (FLAG, amino acid sequence DYKDDDDK) representative TII AVAs to probe lupus nephritis renal tissue. AVAs were then detected using rat IgG anti-FLAG antibodies and an anti-rat IgG tertiary antibody coupled to FITC (**Figure 2a-b**). The distribution of AVA staining was then compared to either V9 or Hoechst stain distributions. Analysis of MPIs (**Figure 2ci**) revealed that both the PB4 and PB5 reversions had diminished intensity colocalized with vimentin. PB4rev also had diminished nuclear colocalization while in PB5, this was increased. When we examined staining co-variance per field of view (FOV), the same trends became more apparent (**Figure 2cii**). Furthermore, visual inspection of representative images was consistent with these results (**Figure 2d**). Probing lupus nephritis biopsies with either PB4mut or PB5mut revealed colocalization with V9 and poor colocalization with Hoechst. In comparison, PB4rev and PB5rev had diminished staining in the vimentin rich areas while PB5rev had increased co-localization with Hoechst. These data demonstrate that both relative binding strength and specificity can be assessed by staining target tissue with tagged AVA IgG monoclonal antibodies.

As HEp-2 cells proved to be a reliable surrogate of tissue, a high-throughput method was developed to determine preferential reactivity of mAbs with subcellular HEp-2 cell domains (**Figure 3**). In addition to providing a simple, unbiased approach to image analysis, we sought to be able to solve complex problems in subcellular localization. For example, it was important to resolve overlapping domains such as when vimentin fibrils extend across the nucleus (Figure 1b).

**Figure 3.**
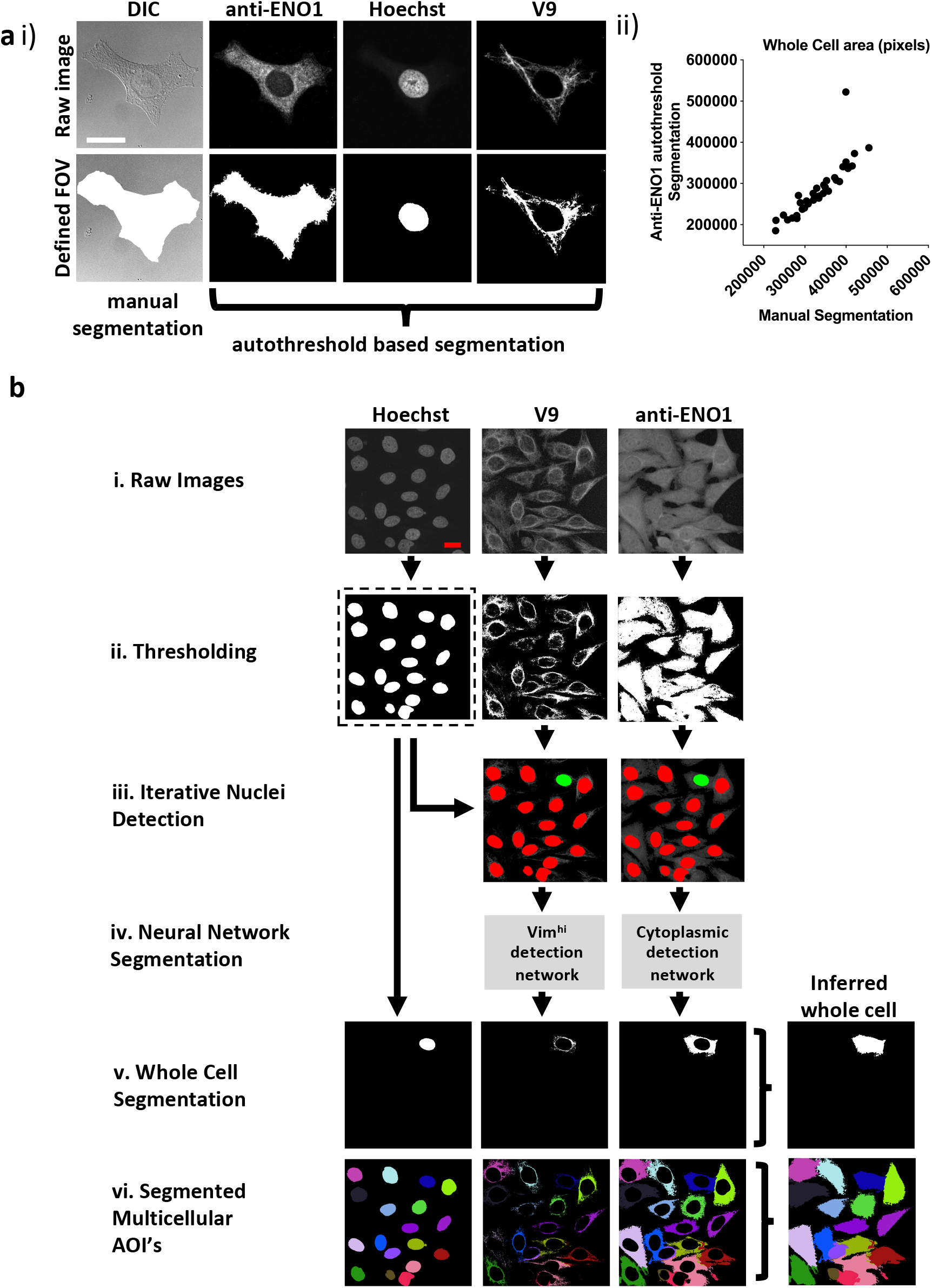
CytoSkaler for automated quantification of subcellular AVA binding. (**a**) Automated whole cell area segmentation was enabled by incorporating an additional fluorescence channel for the anti-a enolase 1 antibody (anti-ENO1) stain, applying autothresholding, despeckling, largest zone selection and subsequent space filling. (**i**) An example of a single cell containing FOV is given. A manually segmented whole cell area is given together with an approximation based on thresholding of the anti-ENO1 fluorescent signal. Vim^hi^ and nuclear regions were segmented by autothresholding as for **Fig 1**. (**ii**) Correlation of autothresholded anti-ENO1 signal to manual segmention of a whole cell area (36 single cell-containing FOVs tested, Spearman r = 0.957 (95% C.I. = 0.915 to 0.978), mean average intersection of unions (IOU) = 0.80. (**b**) Schematic of machine learning based method to train the CytoSkaler program to segment individual cells and respective subcellular areas. (**i**) Acquired FOVs, each containing multiple HEp-2 cells were inputted as respective raw color channels. (**ii**) Channels were autothresholded to yield binary images for each channel based on pixel intensity. (**iii**) Separated nuclear boundaries from the binary Hoechst channel were used as markers to produce RGB images for the V9 and anti-ENO1 channels, yielding a single cell’s nuclear boundary (colored green) and all other cells’ nuclear boundaries colored red. This process was repeated for every nucleus in the Hoechst channel. (**iv**). Two neural networks were trained to separately classify subcellular pixels around Hoechst green boundaries for the V9 and anti-ENO1 channels. Trained on manually segmented cells from 189 ground truth FOVs (2094 cells) with training over 2,300 iterations. These yielded mean IOU scores of 0.96 and 0.89 for the V9 and ENO1 networks, respectively. (**v**) Whole cell segmentations were produced following union of individual channel segmentations and space filling. (**vi**) All segmented cells in a single FOV are marked in a different color to show complete CytoSkaler output of a multicellular zone.

To facilitate automated processing, we stained with an anti-enolase 1 (ENO1) antibody to define the extent of the cytoplasm rather than manual segmentation of DIC images. Therefore, a whole cell was defined as the union of three individually segmented Hoechst, vimentin, and ENO1 regions (**Figure 3ai**). Images were then pre-processed including autothresholding, space filling and selection of largest grouped area to output binary images. Preprocessing was first tested on 36 FOVs that each contained single HEp-2 cells that had been manually segmented (**Figure 3ai**). There was excellent correlation between the automated and hand segmented images as measured by cell size (Spearman’s rho correlation for cell size= 0.957) (**Figure 3aii**) and the intersection over union (IOU) for each whole cell (0.80).

In the new pipeline, raw images (**Figure 3bi**) were first subjected to autothresholding and 2D median filtering (**Figure 3bii**). Next, separated Hoechst boundaries were overlaid onto the V9 and anti-ENO1 filtered images. A separate set of V9 and anti-ENO1 images were created for each Hoechst boundary. In each image, a single Hoechst boundary was colored green and all other Hoechst were colored red. Each nucleus, and ultimately the cellular structures associated with it were considered sequentially (**Figure 3biii**). In contrast to this relatively straightforward segmentation task, close apposition of vimentin fibrils and other cellular domains made segmentation challenging. Therefore, two separate DeepLab v3+ convolutional neural networks (CNNs) were trained to recognize subcellular pixels for the V9 and ENO1 channels (**Figure 3biv**). The learning task of segmenting single cells was simplified from instance segmentation to semantic segmentation by training the models to recognize only a single instance of cellular class instead of segmenting multiple instances of a cellular class in an image. This simplification was facilitated by using colored nuclear boundaries as markers for subcellular pixels. More specifically, the neural networks were trained to recognize a single set of subcellular pixels, associated with a green Hoechst boundary. Next, segmented subcellular areas were combined to create a whole cell segmentation (**Figure 3bv**). This process of color overlay and neural network classification was repeated for all cells in a multicellular image (**Figure 3bvi**). We refer to this analytic pipeline as CytoSkaler. With the two CNNs, CytoSkaler was able to achieve an automated output of single cell segmentation by using multiple subcellular regional stained channels. The mean IOU scores for V9 and ENO1 networks were 0.96 and 0.89, respectively.

We next examined the distribution of somatic hypermutations in PB4 and PB5. In both antibodies, the immunoglobulin light and heavy chains had somatic hypermutations (**Supplemental Figure 2a**). Mutations in both complementary determining regions (CDRs) and framework regions (FRs) were common. To determine which mutated immunoglobulin chain, heavy or light, contributed to vimentin binding, we mixed and matched mut and rev chains and expressed the four resulting iterations for each parent antibody (**Supplemental Figure 2bi**). When these antibodies were tested *in vitro*, it was apparent that the immunoglobulin heavy chain, and not light chain, encoded most vimentin immunoreactivity (**Supplemental Figure 2bii**). As PB5rev conferred nuclear reactivity, commercial anti-dsDNA and anti-histone diagnostic ELISAs were screened for reactivity. This demonstrated that PB5rev conferred histone, and to a lesser degree dsDNA reactivity. Most of this reactivity was provided by the germline immunoglobulin heavy chain (**Supplemental Figure 2biii-iv**).

To better understand the role of SHM in selection for vimentin immunoreactivity, we made a panel of PB5 antibodies in which single encoding V region mutations in the immunoglobulin heavy chain were reverted to predicted germline sequence. When examining the PB5 reversions by ELISA, many appeared to confer increased immunoreactivity with vimentin *in vitro* (**Figure 4a**). In contrast, it was clear that the S36D mutation conferred histone binding. However, only G55V appeared to confer increased colocalization with vimentin fibrils (Vim^hi^) by CytoSkaler (**Figure 4bi**).

**Figure 4.**
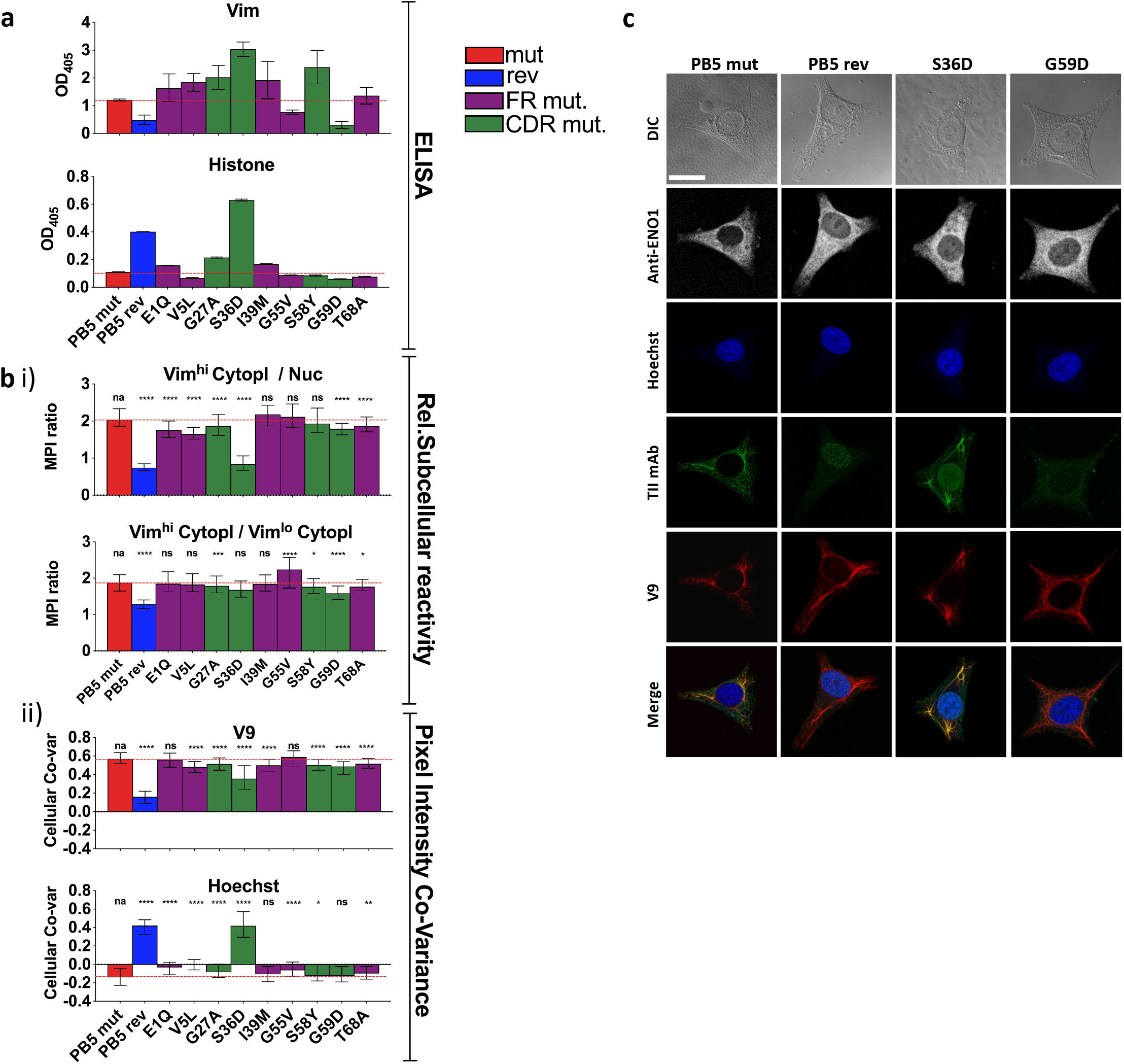
Individual PB5 heavy chain reversions and antigen binding. To determine the relative influences on antigen binding of individual SHMs, variants of TII mAbs PB5 were made such that individual heavy chain SHMs were altered to their predicted germline amino acids. (**a**) mAb reactivity by ELISA. Means and standard deviations are shown. Bar colors represent whether the AVA is fully somatically hypermutated (“mut”), completely reverted to predicted germline (“rev”), or (for a single amino acid reversion) whether the reverted amino lies within framework (FR) region or the complementarity-determining region (CDR). Red dashed lines represent the values for mut AVA binding. (**b-c**) HEp-2 cells were co-stained with V9, Hoechst, anti-ENO1, and one of the indicated PB5 variants. Raw channel data from ten respective FOVs per mAb (approximately ten cells/FOV) was processed using the CytoSkaler. (**bi**) Ratio of MPIs between the indicated subcellular areas medians and interquartile ranges of values for segmented individual cells were plotted. (**bii**) Co-variance of pixel intensities between the respective PB5 variant mAb and V9 or Hoechst. Medians and interquartile ranges of values for segmented individual cells were plotted. (**c**) Single cell examples of the staining patterns yielded by the indicated PB5 variants. White scale bar = 10 microns. Statistical differences (Mann Whitney tests) between the fully mutated AVA and the respective AVA variant is indicated. *p < 0.05, **p < 0.001, ***p < 0.001, ****p < 0.0001 (Mann Whitney).

By CytoSkaler, the PB5 S36D mutation conferred the strongest nuclear reactivity (**Figure 4bii**). This was also evident when representative images were inspected (**Supplemental Figure 3**). Therefore, germline D36 appears to be the main amino acid conferring autoreactivity to the parent antibody. Interestingly, CytoSkaler revealed that the S36D mutation also diminished binding to vimentin. Therefore, selection against the D36 mutation appears to represent selection that prioritizes specificity, and selection against polyreactivity, over affinity for vimentin. In contrast, most other mutations indicate they provide modest increased vimentin binding. These data indicate that CytoSkaler can detect true selection events for specificity. In contrast, one would conclude from the ELISA results that the S36D mutation conferred broad and strong polyreactivity without any specificity. These data provide yet another example in which *in vitro* binding fails to capture specificity that is apparent upon quantitation of *in situ* binding.

We compared the performance of CytoSkaler on subcellular segmentation with CellProfiler, another commonly used imaging tool [24]. The mean IOU scores for V9-Nuc segmentation were 0.9500 and 0.7745 for CytoSkaler and CellProfiler respectively. Analogously, mean IOU scores for ENO1-Nuc segmentation were 0.8306 and 0.7668 for CytoSkaler and CellProfiler respectively (**Supplemental Figure 4**). A developmental version of the CytoSkaler interface is available at https://github.com/awezmm/CytoSkaler.

ELISAs and other *in vitro* binding assays have proven to be profoundly useful in both science and medicine. Furthermore, such simplified systems enable assessments of antibody affinity, direct target binding and epitope mapping. However, our data indicate that for some antibodies ELISAs for polyreactivity over-estimate *in vivo* broad reactivity. This is not necessarily surprising as current polyreactivity assays use highly purified, concentrated and immobilized charged substrates. Such a non-physiological presentation of antigen probably favors nonspecific charge-charge interactions. Therefore, it is almost expected that binding to dsDNA in an ELISA might not always predict binding to genomic DNA in normal cellular or tissue contexts. Our data suggest that the application of confocal imaging and machine learning provides better assessments of antibody specificity than ELISA.

In addition to the better performance on subcellular segmentation, CytoSkaler provides an easy method for calculating mean pixel intensity in any user-defined subcellular region (like Vim^hi^ or Vim^low^) after subcellular segmentation. More specifically, users can define a custom subcellular region that includes the union or subtraction of multiple subcellular regions (e.g. nucleus minus cytoskeleton). Furthermore, CytoSkaler automatically calculates colocalization of antibodies through correlation coefficients between every subcellular channel and the antibody binding channel.

CellProfiler also provides an *IdentifyTertiaryObject* module which can subtract a smaller region from a larger region. However, a smaller region must be completely contained inside a larger region. Furthemore, CellProfiler has no module for combining two separate regions identified at different times. Thus, CellProfiler can be inopportune for analyzing regions that overlap or subsections which involve combination or subtraction of multiple regions. For the purposes of subcellular segmentation and characterization of antibody specificity in user-defined regions through metrics like mean pixel intensity and correlation coefficients, CytoSkaler is more accurate and easier to use than freewares like CellProfiler.

Polyreactivity, as defined in *in vitro* assays, is a common feature of B cell repertoires in both healthy people and those afflicted with lupus [25]. However, it is unclear to what degree this polyreactivity confers risk of *in vivo* autoreactivity. Our data suggest that this risk is less than what has been assumed. Indeed, our results suggest that *in vitro* polyreactivity captures, in some cases, mechanisms that impart *in vivo* binding specificity. To better understand the relationships between autoreactivity, polyreactivity and pathogenicity will require additional approaches, such as that described herein, to better characterize antibody binding specificity *in situ.*

## Supplementary Figure Legends

**Supplemental Figure 1.**
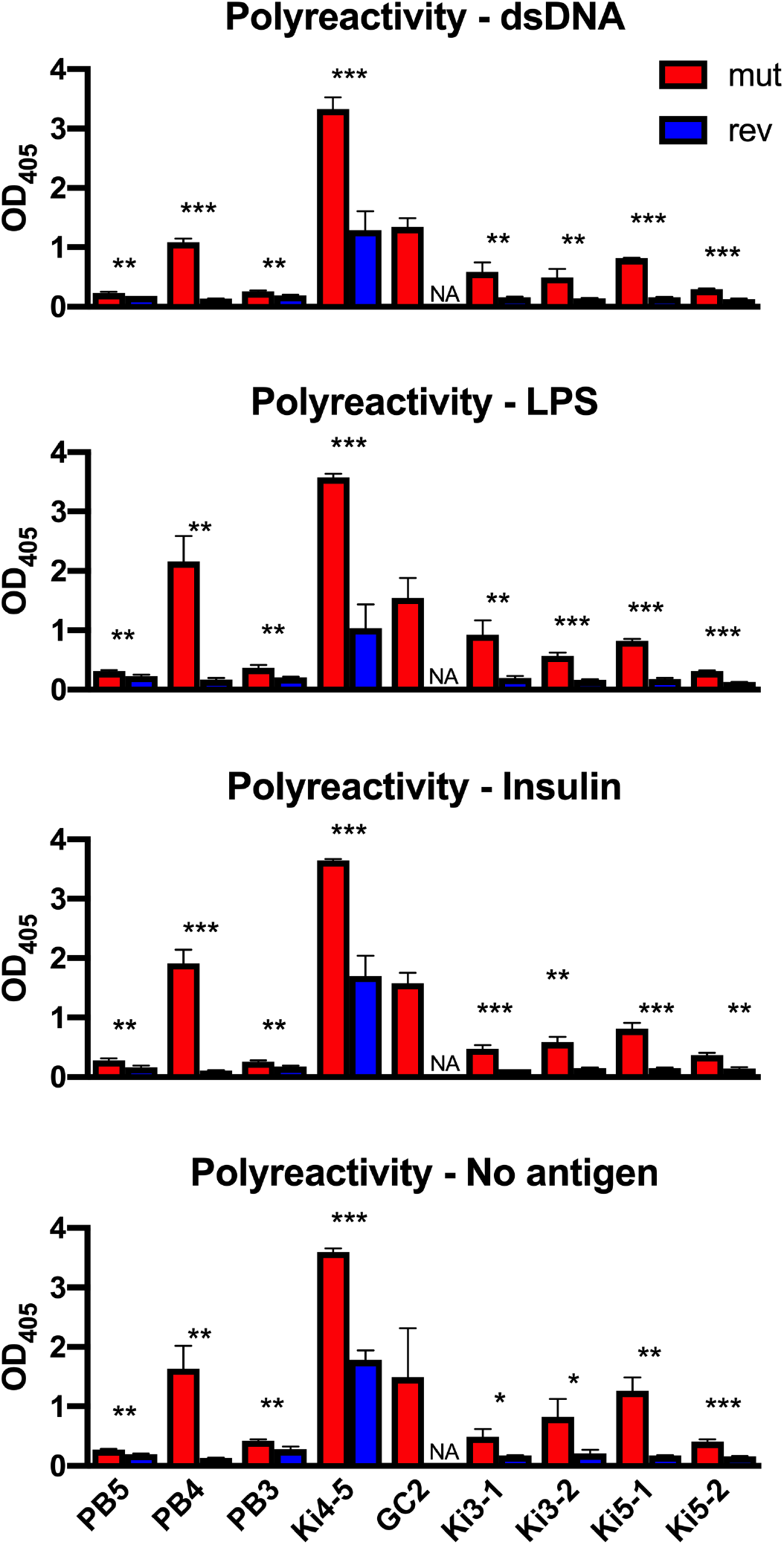
ELISAs for polyreactivity. Reactivity of AVAs (mut and rev) with dsDNA, insulin, LPS, or uncoated plates was measured by ELISA (Raw OD405 values are given and t-tests were performed). *p < 0.05, **p < 0.001, ***p < 0.001.

**Supplemental Figure 2.**
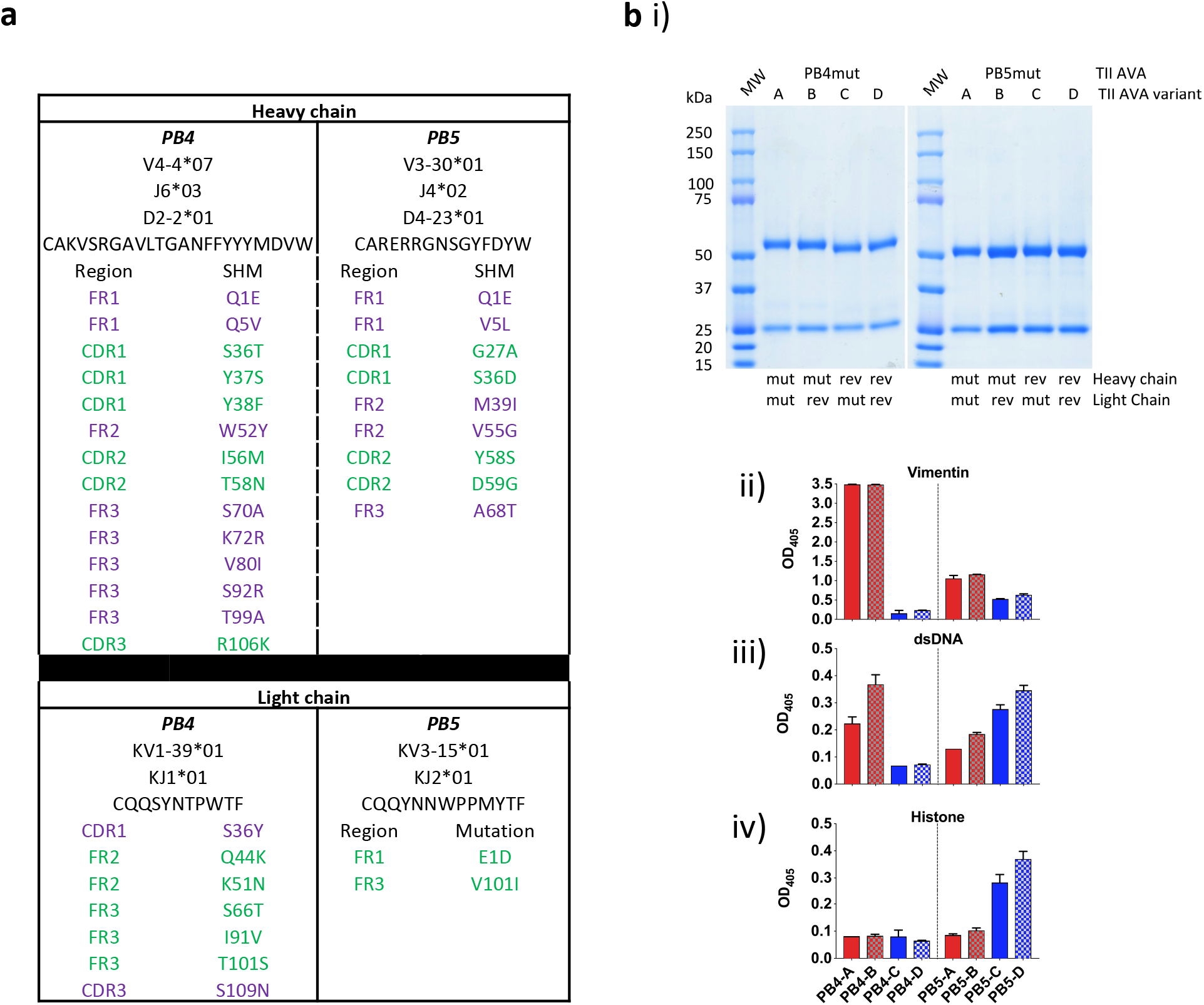
PB4 and PB5 SHMs conferring purified antigen reactivity. (**a**) Somatic hypermutations in TII AVAs PB4mut and PB5mut. The positions of SHMs yielding nonsynonymous mutations are tabulated. Green are in CDRs and purple in FR regions. (**bi**) A representative Coomassie stain of an SDS PAGE gel containing the resolved variants of TII AVAs PB4 and PB5 is displayed. (**bii-iv**) ELISA demonstrating relative reactivities of the respective AVA variants with Vimentin (**ii**) and dsDNA (Inova Diagnostics) (**iii**) and Histone (**iv**).

**Supplemental Figure 3.**
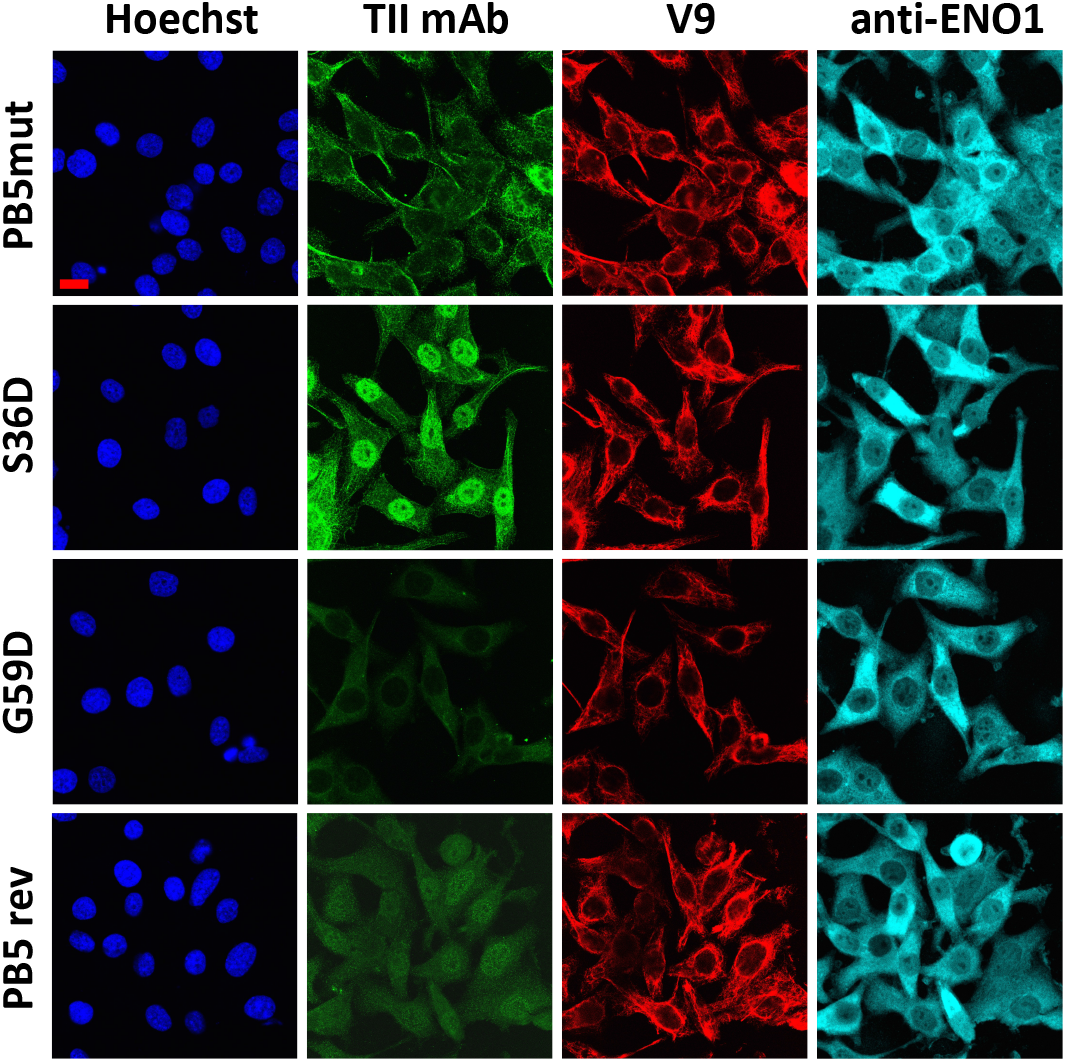
HEp-2 binding by specific PB5 mutants. Examples of FOVs from the indicated AVAs. Representative images from three independent experiments. Red scale bar = 25 microns.

**Supplemental Figure 4.**
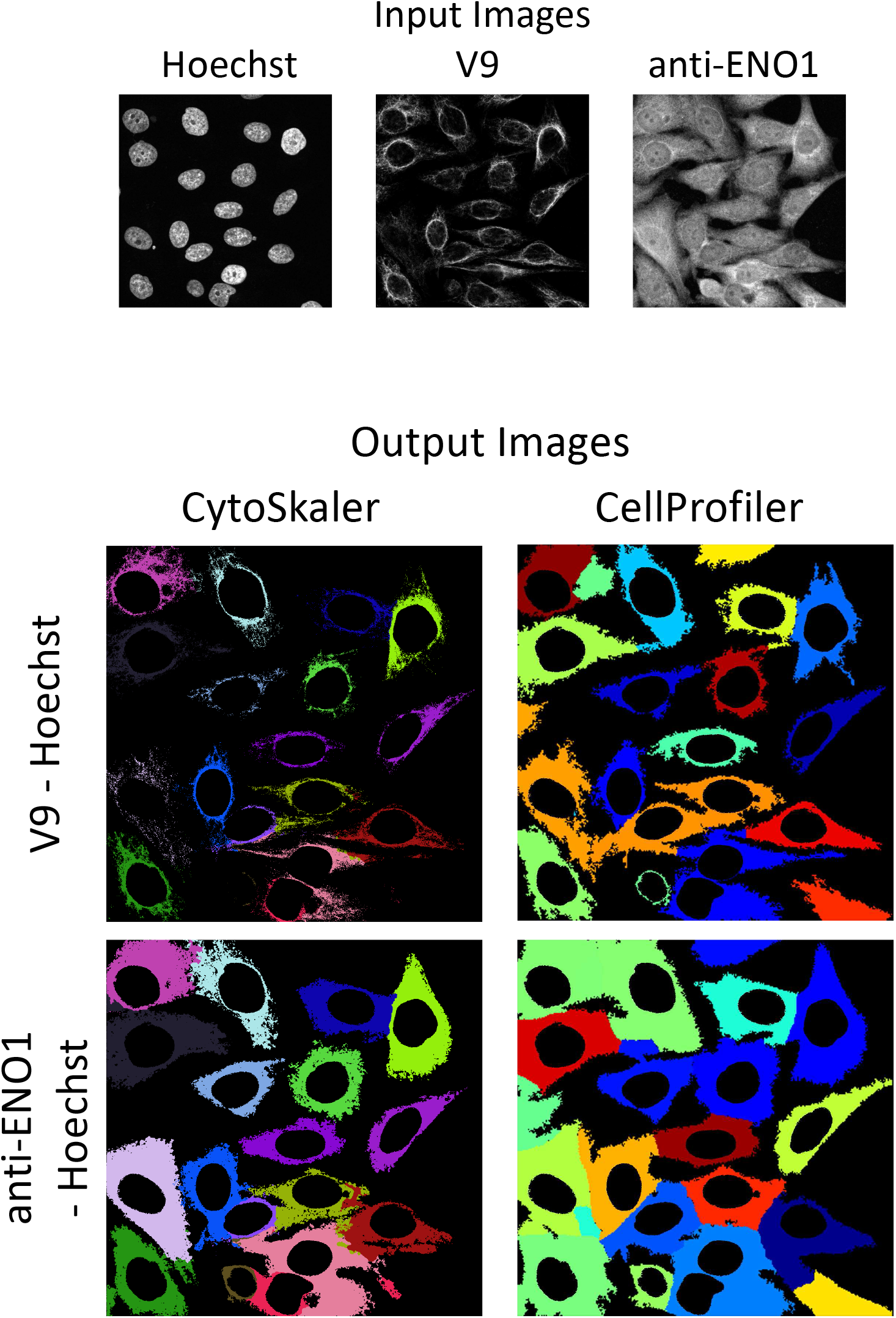
Performance of Cytoskaler compared with CellProfiler. Representative images compare the performance of Cytoskaler and Cell Profiler to perform automated cellular area segmentation of one multicellular HEp-2 FOV. Raw images for respective stains are demonstrated. Bottom panels show automated whole cell segmentation of cells within the FOV using the labelled software. As CellProfiler automatically subtracts the primary region (Hoechst) from the larger secondary regions (V9 and anti-ENO1) for outputs, nuclear regions were subtracted from CytoSkaler outputs as well. The accuracy of the respective software to assign subcellular areas was determined using the IOU metric. CytoSkaler and CellProfiler were both tested on a set of 23 multicellular FOV images, that were not used in CytoSkaler training. The mean IOU scores for “V9 - Hoechst” segmentation were 0.9500 and. 0.7745 for CytoSkaler and CellProfiler respectively. Analogously, mean IOU scores for “anti-ENO1 - Hoechst” segmentation were 0.8306 and 0.7668 for CytoSkaler and CellProfiler respectively.

## REFERENCES

1. Clark, M.R., K. Trotter, and A. Chang, The pathogenesis and therapeutic implications of tubulointerstitial inflammation in human lupus nephritis. Semin Nephrol, 2015. 35(5): p. 455–64.

2. Hsieh, C., et al., Predicting outcomes of lupus nephritis with tubulointerstitial inflammation and scarring. Arthritis Care Res (Hoboken), 2011. 63(6): p. 865–74.

3. Pamfil, C., et al., Intrarenal activation of adaptive immune effectors is associated with tubular damage and impaired renal function in lupus nephritis. Ann Rheum Dis, 2018.

4. Chang, A., et al., In situ B cell-mediated immune responses and tubulointerstitial inflammation in human lupus nephritis. J Immunol, 2011. 186(3): p. 1849–60.

5. Shen, Y., et al., Association of intrarenal B-cell infiltrates with clinical outcome in lupus nephritis: a study of 192 cases. Clin Dev Immunol, 2012. 2012: p. 967584.

6. Liarski, V.M., et al., Cell distance mapping identifies functional T follicular helper cells in inflamed human renal tissue. Sci Transl Med, 2014. 6(230): p. 230ra46.

7. Kinloch, A.J., et al., Vimentin is a dominant target of in situ humoral immunity in human lupus tubulointerstitial nephritis. Arthritis Rheumatol, 2014. 66(12): p. 3359–70.

8. Kinloch, A.J., et al., Serum Anti-Vimentin Autoantibodies Predict Loss of Renal Function in Lupus Nephritis under revision, 2020.

9. Bilalic, S., et al., Lymphocyte activation induces surface expression of an immunogenic vimentin isoform. Transpl Immunol, 2012. 27: p. 101–106.

10. Mor-Vaknin, N., et al., Vimentin is secreted by activated macrophages. Nat Cell Biol, 2003. 5: p. 59–63.

11. Tan, E.M., et al., The 1982 revised criteria for the classification of systemic lupus erythematosus. Arthritis Rheum, 1982. 25(11): p. 1271–7.

12. Wellmann, U., et al., The evolution of human anti-double-stranded DNA autoantibodies. Proc Natl Acad Sci U S A, 2005. 102(26): p. 9258–63.

13. Guo, W., et al., Somatic hypermutation as a generator of antinuclear antibodies in a murine model of systemic autoimmunity. J Exp Med, 2010. 207(10): p. 2225–37.

14. Shlomchik, M.J., et al., The role of clonal selection and somatic mutation in autoimmunity. Nature, 1987. 328(6133): p. 805–11.

15. Wardemann, H., et al., Predominant autoantibody production by early human B cell precursors. Science, 2003. 301(5638): p. 1374–7.

16. Yurasov, S., et al., Defective B cell tolerance checkpoints in systemic lupus erythematosus. J Exp Med, 2005. 201: p. 703–711.

17. Smith, K., et al., Rapid generation of fully human monoclonal antibodies specific to a vaccinating antigen. Nat Protoc, 2009. 4(3): p. 372–84.

18. Kaur, K., et al., High Affinity Antibodies against Influenza Characterize the Plasmablast Response in SLE Patients After Vaccination. PLoS One, 2015. 10(5): p. e0125618.

19. Kinloch, A.J., et al., In Situ Humoral Immunity to Vimentin in HLA-DRB1‪03(+) Patients With Pulmonary Sarcoidosis. Front Immunol, 2018. 9: p. 1516.

20. Henry, C., et al., Influenza Virus Vaccination Elicits Poorly Adapted B Cell Responses in Elderly Individuals. Cell Host Microbe, 2019. 25(3): p. 357–366 e6.

21. Bolte, S. and F.P. Cordelieres, A guided tour into subcellular colocalization analysis in light microscopy. J Microsc, 2006. 224(Pt 3): p. 213–32.

22. Andrews, S.F., et al., Immune History Profoundly Affects Broadly Protective B Cell Responses to Influenza. Sci Transl Med, 2015. 7: p. 316ra192.

23. Herrmann, H., et al., Intermediate filaments: primary determinants of cell architecture and plasticity. J Clin Invest, 2009. 119(7): p. 1772–83.

24. Kamentsky, L., et al., Improved structure, function and compatibility for CellProfiler: modular high-throughput image analysis software. Bioinformatics, 2011. 27(8): p. 1179–80.

25. Yurasov, S., et al., Defective B cell tolerance checkpoints in systemic lupus erythematosus. J Exp Med, 2005. 201(5): p. 703–11.

