## Supplemental Table 1 for "*In situ* humoral selection in human lupus tubulointerstitial inflammation"

| **Supplemental Table 1: nucleotide sequences** | | | |
| --- | --- | --- | --- |
| **Type** | **name** | **vector** | **mAb / insert or primer sequence** |
| ***mAbs*** |  |  |  |
| TII mAb H chain | GC2H | AbVec-hIgG1 | [7] |
| TII mAb H chain | Ki3-1H | AbVec-hIgG1 | [7] |
| TII mAb H chain | Ki3-2H | AbVec-hIgG1 | [7] |
| TII mAb H chain | Ki5-1H | AbVec-hIgG1 | [7] |
| TII mAb H chain | Ki5-2H | AbVec-hIgG1 | [7] |
| TII mAb H chain | Ki4-5H | AbVec-hIgG1 | [7] |
| TII mAb H chain | PB3H | AbVec-hIgG1 | [7] |
| TII mAb H chain | PB4H | AbVec-hIgG1 | [7] |
| TII mAb H chain | PB5H | AbVec-hIgG1 | [7] |
| TII mAb H chain rev | GC2Hrev | AbVec-hIgG1 | CTGCAACCGGTGTACATTCTgaggtgcagctggtggagactggaggaggcttgatccagcctggggggtccctgagactctcctg  TgcagcctctgggttcaccgtcagtAgcaactacatgagctgggtccgccaggctccagggaaggggctggagtgggtctcagttatttat  agcggtggtagcacatactacgca  gactccgtgaagggccgattcaccatctccagagacaattccaagaacacgctgtatcttcaaatgaacagcctgagagccgaggacacg  gccgtgtattactgtgcgagagagccctggttgggtcatttgggggagtactactttgactactggggccagggaaccctggtcaccgtc  tcctcagCGTCGACTTCGCA |
| TII mAb H chain rev | Ki3-1Hrev | AbVec-hIgG1 | CTGCAACCGGTGTACATTCTgaggtgcagctggtggagtctgggggaggcttggtccagcctggggggtccctgagactctcctgt  gcagcctctggattcacctttagt  agctattggatgagctgggtccgccaggctccagggaaggggctggagtgggtggccaacataaagcaagatggaagtgagaaatactat  gtggactctgtgaagggccgattcaccatctccagagacaacgccaagaactcactgtatctgcaaatgaacagcctgagagccgaggac  acggctgtgtattactgtgcgagagaattactatggttcggggactactttgactactggggccagggaaccctggtcaccgtctcctca  gCGTCGACTTCGCA |
| TII mAb H chain rev | Ki3-2Hrev | AbVec-hIgG1 | CTGCAACCGGTGTACATTCTgaggtgcagctggtggagtctgggggaggcttggtccagcctggggggtccctgagactctcctgtg  cagcctctggattcacctttagt  agctattggatgagctgggtccgccaggctccagggaaggggctggagtgggtggccaacataaagcaagatggaagtgagaaatactat  gtggactctgtgaagggccgattcaccatctccagagacaacgccaagaactcactgtatctgcaaatgaacagcctgagagccgaggac  acggctgtgtattactgtgcgagagaattactatggttcggggactactttgactactggggccagggaaccctggtcaccgtctcctca  gCGTCGACTTCGCA |
| TII mAb H chain rev | Ki5-1Hrev | AbVec-hIgG1 | CTGCAACCGGTGTACATTCTgaggtgcagctggtggagtctgggggaggcttggtaaagcctggggggtcccttagactctcct  gtgcag  cctctggattcactttcagtaacgcctggatgagctgggtccgccaggctccagggaaggggctggagtgggttggccgtattaaaagca  aaactgatggtgggacaacagactacgctgcacccgtgaaaggcagattcaccatctcaagagatgattcaaaaaacacgctgtatctgc  aaatgaacagcctgaaaaccgaggacacagccgtgtattactgtaccacagaggacgcaggggaagatgatgcttttgatatctggggcc  aagggacaatggtcaccgtctcttcagCGTCGACCAAGGG |
| TII mAb H chain rev | Ki5-2Hrev | AbVec-hIgG1 | CTGCAACCGGTGTACATTCTcaggtgcagctgcaggagtcgggcccaggactggtgaagccttcggagaccctgtccctcacc  tgcactg  tctctggttactccatcagcagtggttactactggggctggatccggcagcccccagggaaggggctggagtggattgggagtatctatc  atagtgggagcacctactacaacccgtccctcaagagtcgagtcaccatatcagtagacacgtccaagaaccagttctccctgaagctga  gctctgtgaccgccgcagacacggccgtgtattactgtgcgagagacgggacagcggctggtgtatggggggactactttgactactggg  gccagggaaccctggtcaccgtctcctcagCGTCGACCAAGGG |
| TII mAb H chain rev | Ki4-5Hrev | AbVec-hIgG1 | CTGCAACCGGTGTACATTCTcaggtgcagctgcaggagtcgggcccaggactggtgaagccttcacagaccctgtccctcacc  tgcactgtctctggtggctccatcagc  agtggtagttactactggagctggatccgccagcacccagggaagggcctggagtggattgggtacatctattacagtgggagcacctac  tacaacccgtccctcaagagtcgagttaccatatcagtagacacgtctaagaaccagttctccctgaagctgagctctgtgactgccgcg  gacacggccgtgtattactgtgcgaggtacgcggggcgcagcagctggtaccaaaccgggtacacctccatgtcctactactttgactac  tggggccagggaaccctggtcaccgtctcctcagCGTCGACTTCGCA |
| TII mAb H chain rev | PB3Hrev | AbVec-hIgG1 | CTGCAACCGGTGTACATTCTcaggtgcagctggtggagtctgggggaggcgtggtccagcctgggaggtccctgagactctcctg  tgcagcctctggattcaccttcagt  agctatgctatgcactgggtccgccaggctccaggcaaggggctggagtgggtggcagttatatcatatgatggaagcaataaatactac  gcagactccgtgaagggccgattcaccatctccagagacaattccaagaacacgctgtatctgcaaatgaacagcctgagagctgaggac  acggctgtgtattactgtgcgagagatcccgggggttatggggccctggacaactggttcgacccctggggccagggaaccctggtcacc  gtctcctcagCGTCGACTTCGCA |
| TII mAb H chain rev | PB4Hrev | AbVec-hIgG1  &  AbVec-hIgG1-DYKDDDDK | CTGCAACCGGTGTACATTCTcaggtgcagctgcaggagtcgggcccaggactggtgaagccttcggagaccctgtccctcacctg  cactgtctctggtggctccatcagt  agttactactggagctggatccggcagcccgccgggaagggactggagtggattgggcgtatctataccagtgggagcaccaactacaac  ccctccctcaagagtcgagtcaccatgtcagtagacacgtccaagaaccagttctccctgaagctgagctctgtgaccgccgcggacacg  gccgtgtattactgtgcgagagagtctcgtggcgcagtgttaacaggtgattactactactactactacatggacgtctggggcaaaggg  accacggtcaccgtctcctcagCGTCGACTTCGCA |
| TII mAb H chain rev | PB5Hrev | AbVec-hIgG1  &  AbVec-hIgG1-DYKDDDDK | CTGCAACCGGTGTACATTCTcaggtgcagctggtggagtctgggggaggcgtggtccagcctgggaggtccctgagactctcctg  tgcagcctctggattcaccttcagt  agctatgctatgcactgggtccgccaggctccaggcaaggggctggagtgggtggcagttatatcatatgatggaagcaataaatactac  gcagactccgtgaagggccgattcaccatctccagagacaattccaagaacacgctgtatctgcaaatgaacagcctgagagctgaggac  acggctgtgtattactgtgcgagagagaggcgaggtaactccgactactttgactactggggccagggaaccctggtcaccgtctcctca  gCGTCGACTTCGCA |
| TII mAb L chain | GC2L | K | [7] |
| TII mAb L chain | Ki3-1 L | λ | [7] |
| TII mAb L chain | Ki3-2 L | λ | [7] |
| TII mAb L chain | Ki5-1 L | λ | [7] |
| TII mAb L chain | Ki5-2 L | λ | [7] |
| TII mAb L chain | Ki4-5 L | K | [7] |
| TII mAb L chain | PB3 L | K | [7] |
| TII mAb L chain | PB4 L | K | [7] |
| TII mAb L chain | PB5 L | K | [7] |
| TII mAb L chain rev | GC2 Lrev | K | Tacattcggacatcgtgatgacccagtctccagactccctggctgtgtctctgggcgagagggccaccat  Caactgcaagtccagccagagtgttttatacagctccaacaataagaactacttagcttggtaccagcag  Aaaccaggacagcctcctaagctgctcatttactgggcatctacccgggaatccggggtccctgaccgattcagtggcagcgggtctg  ggacagatttcactctca  ccatcagcagcctgcaggctgaagatgtggcagtttattactgtcagcaatattatagtttcggccaagggaccaaggtggaaatcaa  acgtacgtggctgc |
| TII mAb L chain rev | Ki3-1 Lrev | λ | Ctgctaccggttcttgggcccagtctgccctgactcagcctgcctccgtgtctgggtctcctggacagtcgatcaccatctcctgcactgg  aaccagcagtgacgttggt  ggttataactatgtctcctggtaccaacagcacccaggcaaagcccccaaactcatgatttatgaggtcagtaatcggccctcaggggtt  tctaatcgcttctctggctccaagtctggcaacacggcctccctgaccatctctgggctccaggctgaggacgaggctgattattactgc  agctcatatacaagcagcagcactctcgtattcggcggagggaccaagctgaccgtcctaggtcagcccaaggctgccccctcggtcact  ctgttcccACCCTCGAGTGAGGAG |
| TII mAb L chain rev | Ki3-2 Lrev | λ | CTGCTACCGGTTCCTGGGCCcagtctgccctgactcagcctccctccgcgtccgggtctcctggacagtcagtcaccatctcctg  cactggaaccagcagtgacgttggt  ggttataactatgtctcctggtaccaacagcacccaggcaaagcccccaaactcatgatttatgaggtcagtaagcggccctcaggggtc  cctgatcgcttctctggctccaagtctggcaacacggcctccctgaccgtctctgggctccaggctgaggatgaggctgattattactgc  agctcatatgcaggcagcaacaatttggtattcggcggagggaccaagctgaccgtcctaggtcagcccaaggctgccccctcggtcact  ctgttcccACCCTCGAGTGAGGAG |
| TII mAb L chain rev | Ki5-1 Lrev | λ | CTGCAACCGGTGTACATTCTcagtctgccctgactcagcctgcctccgtgtctgggtctcctggacagtcgatcaccatctcctgc  actggaaccagcagtgacgttggtggttataactatgtctcctggtaccaacagcacccaggcaaagcccccaaactcatgatttatgaggtca  gtaatcggccctcaggggtttctaatcgcttctctggctccaagtctggcaacacggcctccctgaccatctctgggctccaggctgagg  acgaggctgattattactgcagctcatatacaagcagcagcactctcgtcttcggaactgggaccaaggtcaccgtcctaggtcagccca  aggccaaccccactgtcactctgttcccgccCTCGAGTGAGGAG |
| TII mAb L chain rev | Ki5-2 Lrev | λ | CTGCAACCGGTGTACATTCTcagtctgccctgactcagcctgcctccgtgtctgggtctcctggacagtcgatcaccatctcctgcactg  gaaccagcagtgacgttggtggttataactatgtctcctggtaccaacagcacccaggcaaagcccccaaactcatgatttatgaggtca  gtaatcggccctcaggggtttctaatcgcttctctggctccaagtctggcaacacggcctccctgaccatctctgggctccaggctgagg  acgaggctgattattactgcagctcatatacaagcagcagcactctcgtcttcggaactgggaccaaggtcaccgtcctaggtcagccca  aggccaaccccactgtcactctgttcccgccCTCGAGTGAGGAG |
| TII mAb L chain rev | Ki4-5 Lrev | K | CTGCAACCGGTGTACATTCGgacatccagatgacccagtctccttccaccctgtctgcatctgtaggagacagagtcaccatcac  ttgccgggccagtcagagtattagt  agctggttggcctggtatcagcagaaaccagggaaagcccctaagctcctgatctataaggcgtctagtttagaaagtggggtcccatca  aggttcagcggcagtggatctgggacagaattcactctcaccatcagcagcctgcagcctgatgattttgcaacttattactgccaacag  tataatagttattctcccactttcggccctgggaccaaagtggatatcaaaCGTACGGTGGC |
| TII mAb L chain rev | PB3 Lrev | K | CTGCAACCGGTGTACATTCTgacatccagatgacccagtctccatcctccctgtctgcatctgtaggagacagagtcaccatcac  ttgccgggcaagtcagagcattagc  agctatttaaattggtatcagcagaaaccagggaaagcccctaagctcctgatctatgctgcatccagtttgcaaagtggggtcccatca  aggttcagtggcagtggatctgggacagatttcactctcaccatcagcagtctgcaacctgaagattttgcaacttactactgtcaacag  agttacagtacccctccgacgttcggccaagggaccaaggtggaaatcaaaCGTACGGTGGC |
| TII mAb L chain rev | PB4 Lrev | K | CTGCAACCGGTGTACATTCTgacatccagatgacccagtctccatcctccctgtctgcatctgtaggagacagagtcaccatc  acttgccgggcaagtcagagcattagc  agctatttaaattggtatcagcagaaaccagggaaagcccctaagctcctgatctatgctgcatccagtttgcaaagtggggtcccatca  aggttcagtggcagtggatctgggacagatttcactctcaccatcagcagtctgcaacctgaagattttgcaacttactactgtcaacag  agttacagtacccctccgacgttcggccaagggaccaaggtggaaatcAAACGTACGGTGGC |
| TII mAb L chain rev | PB5Lrev | K | CTGCAACCGGTGTACATGGGgaaatagtgatgacgcagtctccagccaccctgtctgtgtctccaggggaaagagccaccct  ctcctgcagggccagtcagagtgttagc  agcaacttagcctggtaccagcagaaacctggccaggctcccaggctcctcatctatggtgcatccaccagggccactggtatcccagcc  aggttcagtggcagtgggtctgggacagagttcactctcaccatcagcagcctgcagtctgaagattttgcagtttattactgtcagcag  tataataactggcctcccatgtacacttttggccaggggaccaagctggagatcAAACGTACGGTGGC |
| PB5H single a.a rev | PB5HE1Q | AbVec-hIgG1-DYKDDDDK | CTGCAACCGGTGTACATTCTcaggtgcagctgttggagtctggggggggcgtggtccagcctgggaggtccctgagactc tcctgtgcagcctctgcattcaccttcagtgactatgctatccactgggtccgccaggct ccaggcaaggggctggagtgggtggcaggtatatcatctggtggaagcaataaatactac acagactccgtgaagggccgattcaccatctccagagacaattccaagaacacactctat ctccaaatgaacagcctgagagctgaggacacagctgtctattactgtgcgagagagagg cgaggtaactccggctactttgactactggggccagggaaccctggtcaccgtctcctcagCGTCGACTTCGCA |
| PB5H single a.a rev | PB5HV5L | AbVec-hIgG1-DYKDDDDK | CTGCAACCGGTGTACATTCTgaggtgcagctggtggagtctggggggggcgtggtccagcctgggaggtccctgagactc tcctgtgcagcctctgcattcaccttcagtgactatgctatccactgggtccgccaggct ccaggcaaggggctggagtgggtggcaggtatatcatctggtggaagcaataaatactac acagactccgtgaagggccgattcaccatctccagagacaattccaagaacacactctat ctccaaatgaacagcctgagagctgaggacacagctgtctattactgtgcgagagagagg cgaggtaactccggctactttgactactggggccagggaaccctggtcaccgtctcctcagCGTCGACTTCGCA |
| PB5H single a.a rev | PB5HG27A | AbVec-hIgG1-DYKDDDDK | CTGCAACCGGTGTACATTCTgaggtgcagctgttggagtctggggggggcgtggtccagcctgggaggtccctgagactc tcctgtgcagcctctggattcaccttcagtgactatgctatccactgggtccgccaggct ccaggcaaggggctggagtgggtggcaggtatatcatctggtggaagcaataaatactac acagactccgtgaagggccgattcaccatctccagagacaattccaagaacacactctat ctccaaatgaacagcctgagagctgaggacacagctgtctattactgtgcgagagagagg cgaggtaactccggctactttgactactggggccagggaaccctggtcaccgtctcctcagCGTCGACTTCGCA |
| PB5H single a.a rev | PB5HS36D | AbVec-hIgG1-DYKDDDK | CTGCAACCGGTGTACATTCTgaggtgcagctgttggagtctggggggggcgtggtccagcctgggaggtccctgagactc tcctgtgcagcctctgcattcaccttcagtagctatgctatccactgggtccgccaggct ccaggcaaggggctggagtgggtggcaggtatatcatctggtggaagcaataaatactac acagactccgtgaagggccgattcaccatctccagagacaattccaagaacacactctat ctccaaatgaacagcctgagagctgaggacacagctgtctattactgtgcgagagagagg cgaggtaactccggctactttgactactggggccagggaaccctggtcaccgtctcctcagCGTCGACTTCGCA |
| PB5H single a.a rev | PB5HI39M | AbVec-hIgG1-DYKDDDK | CTGCAACCGGTGTACATTCTgaggtgcagctgttggagtctggggggggcgtggtccagcctgggaggtccctgagactc tcctgtgcagcctctgcattcaccttcagtgactatgctatgcactgggtccgccaggct ccaggcaaggggctggagtgggtggcaggtatatcatctggtggaagcaataaatactac acagactccgtgaagggccgattcaccatctccagagacaattccaagaacacactctat ctccaaatgaacagcctgagagctgaggacacagctgtctattactgtgcgagagagagg cgaggtaactccggctactttgactactggggccagggaaccctggtcaccgtctcctcagCGTCGACTTCGCA |
| PB5H single a.a rev | PB5HG55V | AbVec-hIgG1-DYKDDDK | CTGCAACCGGTGTACATTCTgaggtgcagctgttggagtctggggggggcgtggtccagcctgggaggtccctgagactc tcctgtgcagcctctgcattcaccttcagtgactatgctatccactgggtccgccaggct ccaggcaaggggctggagtgggtggcagttatatcatctggtggaagcaataaatactac acagactccgtgaagggccgattcaccatctccagagacaattccaagaacacactctat ctccaaatgaacagcctgagagctgaggacacagctgtctattactgtgcgagagagagg cgaggtaactccggctactttgactactggggccagggaaccctggtcaccgtctcctcagCGTCGACTTCGCA |
| PB5H single a.a rev | PB5HS58Y | AbVec-hIgG1-DYKDDDK | CTGCAACCGGTGTACATTCTgaggtgcagctgttggagtctggggggggcgtggtccagcctgggaggtccctgagactc tcctgtgcagcctctgcattcaccttcagtgactatgctatccactgggtccgccaggct ccaggcaaggggctggagtgggtggcaggtatatcatatggtggaagcaataaatactac acagactccgtgaagggccgattcaccatctccagagacaattccaagaacacactctat ctccaaatgaacagcctgagagctgaggacacagctgtctattactgtgcgagagagagg cgaggtaactccggctactttgactactggggccagggaaccctggtcaccgtctcctcagCGTCGACTTCGCA |
| PB5H single a.a rev | PB5HG59D | AbVec-hIgG1-DYKDDDK | CTGCAACCGGTGTACATTCTgaggtgcagctgttggagtctggggggggcgtggtccagcctgggaggtccctgagactc tcctgtgcagcctctgcattcaccttcagtgactatgctatccactgggtccgccaggct ccaggcaaggggctggagtgggtggcaggtatatcatctgatggaagcaataaatactac acagactccgtgaagggccgattcaccatctccagagacaattccaagaacacactctat ctccaaatgaacagcctgagagctgaggacacagctgtctattactgtgcgagagagagg cgaggtaactccggctactttgactactggggccagggaaccctggtcaccgtctcctcagCGTCGACTTCGCA |
| PB5H single a.a rev | PB5HT68A | AbVec-hIgG1-DYKDDDK | CTGCAACCGGTGTACATTCTgaggtgcagctgttggagtctggggggggcgtggtccagcctgggaggtccctgagactc tcctgtgcagcctctgcattcaccttcagtgactatgctatccactgggtccgccaggct ccaggcaaggggctggagtgggtggcaggtatatcatctggtggaagcaataaatactac gcagactccgtgaagggccgattcaccatctccagagacaattccaagaacacactctat ctccaaatgaacagcctgagagctgaggacacagctgtctattactgtgcgagagagagg cgaggtaactccggctactttgactactggggccagggaaccctggtcaccgtctcctcagCGTCGACTTCGCA |
| FLAG insert | DYKDDDDK H-chain 3’ plus | pUC57 | Ggtgtacaccctgcccccatcccgggatgagctgaccaagaaccaggtcagcctgacctgcctggtcaaaggcttctatcccagcgacat  Cgccgtggagtgggagagcaatgggcagccggagaacaactacaagaccacgcctcccgtgctggactccgacggctccttcttcctct  Acagcaagctcaccgtggacaagagcaggtggcagcaggggaacgtcttctcatgctccgtgatgcatgaggctctgcacaaccactac  Acgcagaagagcctctccctgtctccgggtaaagactacaaagacgatgacgacaagtgaagcttggccgccatggcccaacttgtttat  Tgcagcttataatggttacaaataaagcaatagcatcacaaatttcacaaataaagcatttttttcactgcattctagttgtggtttgtccaa  Actcatcaatgtatcttatcatgtctggatcgatcgggaattaattcggcgcagcaccatggcctgaaataacctctgaaagaggaacttg  gttaggtaccttctgaggcggaaagaaccagctgtggaatgt |
| Primers | IgG INT 2 | NA | GTGCTGGACTCCGACGGCTCCTTC |
