## Supplemental Table 2 for "*In situ* humoral selection in human lupus tubulointerstitial inflammation"

| **Supplemental Table 2: KDs of TII AVAs** | |
| --- | --- |
| *mAb* | *Kd* |
| PB4 | 3.578E-08 |
| PB4rev | low |
| PB5 | 5.06333E-07 |
| PB5rev | 8.67667E-07 |
| Ki3-1 | 2.846E-07 |
| Ki3-1rev | 5.358E-07 |
| Ki3-2 | 9.652E-08 |
| Ki3-2rev | 1.974E-07 |
| Ki4-5 | 6.817E-08 |
| Ki4-5rev | 2.1005E-07 |
| Ki5-1 | 5.349E-07 |
| Ki5-1rev | low |
| Ki5-2 | 4.208E-07 |
| Ki5-2rev | low |
| PB3 | 0.000001059 |
| PB3rev | low |
| GC2 | 0.000002246 |
| “low”=mAbs with too little reactivity for Kd calculation | |
